# Topological Tuning of DNA Mobility in Entangled Solutions of Supercoiled Plasmids

**DOI:** 10.1101/2020.09.21.306092

**Authors:** Jan Smrek, Jonathan Garamella, Rae Robertson-Anderson, Davide Michieletto

## Abstract

Understanding the behaviour of ring polymers in dense solutions is one of the most intriguing problems in polymer physics with far-reaching implications from material science to genome biology. Thanks to its natural occurrence in circular form, DNA has been intensively employed as a proxy to study the fundamental physics of ring polymers in different topological states. Yet, torsionally constrained – such as supercoiled – topologies have been largely neglected so far. Extreme entanglement and high supercoiling levels are commonly found in the genetic material of both pro- and eukaryotes and, at the same time, the applicability of existing theoretical models to dense supercoiled DNA is unknown. To address this gap, here we couple large scale Molecular Dynamics (MD) simulations of twistable chains together with Differential Dynamic Microscopy (DDM) of entangled supercoiled DNA plasmids. We discover that, strikingly, and contrarily to what is generally assumed in the literature, a higher degree of supercoiling increases the average size of plasmids in entangled solutions. At the same time, we discover that this is accompanied by an unexpected enhancement in DNA mobility. We reconcile these apparently contradicting findings as due to the fact that supercoiling drives highly asymmetric plasmid conformations, decreases inter-plasmids entanglements and, in particular, reduces the number of threadings between DNA rings. Our numerical and experimental results also suggest a way to topologically tune DNA mobility via supercoiling, thus enabling the orthogonal control over the (micro)rheology of DNA-based complex fluids with respect to other traditional methods such as DNA length or concentration.

## 1 Introduction

The deoxyribonucleic acid (DNA) is not only the central molecule of life but it is now increasingly employed for bio-compatible and responsive materials – such as DNA hydrogels (*1*) and origami (*2*) – with applications in medicine and nanotechnology (*3*). One feature that renders DNA a unique polymer is its ability to encode information, and this is now extensively leveraged to make complex structures (*3–5*) and even self-replicating materials (*6*); another feature that distinguishes DNA from other synthetic polymers is its unique geometry, i.e. that of a (righthanded) helix with a well-defined pitch, which entails that DNA can display both bending and torsional stiffness (*7*). Oppositely to the information-encoding feature of DNA, its geometrical features are far less exploited to create synthetic materials. In fact, DNA is at present largely employed to make up biopolymer complex fluids in its simplest geometrical forms, i.e. that of a linear or relaxed circular (torsionally unconstrained) molecule (*8–12*). In spite of this, most naturally occurring DNA is under torsional and topological constraints, either because it is circular and non-nicked, as in bacteria, or because of the binding of proteins that restrict the relative rotation of base-pairs, as in eukaryotes (*13–16*). The torsional stress stored in a closed DNA molecule cannot be mechanically relaxed (in absence of Topoisomerase proteins) but only re-arranged or converted into bending in order to minimise the overall conformational free energy (*17–19*). This entails that supercoiling – the linking deficit between sister DNA strands with respect to their relaxed state – can carry conformational information (*20*) which can affect the static and dynamic properties of DNA plasmids (*17*) and even regulate gene transcription (*21*). Here, we propose that supercoiling may also be leveraged to tune the dynamics of DNA plasmids in solution, thus potentially allowing for fine control over the rheology of DNA-based complex fluids in a way that is orthogonal to varying DNA length (*22*), concentration (*23*) or architecture (*9, 24*). Finally, entangled solutions of DNA plasmids are not only interesting because of potential applications in bio and nanotechnology but also as enabling us to study fundamental questions on the physics of ring polymers – one of the most active fields of soft matter research (*25–32*) – thanks to the extremely precise control over DNA lengths and topology (*9–11*) and access to sophisticated visualisation techniques (*33*).

To characterise the effect of DNA supercoiling on the rheology of entangled solutions of plasmids, here we perform large scale Molecular Dynamics simulations of entangled DNA plasmids (Fig. 1A-C), modelled as coarse-grained twistable chains (*34*). We discover that while isolated DNA plasmids typically display a collapse with increasing levels of supercoiling (estimated via simulations (*35*) or gel electrophoresis (*36*)), here we show that entangled DNA plasmids typically increase their average size with supercoiling. Importantly, we further discover that in spite of this swelling, larger supercoiling is accompanied by an *enhanced* mobility of the plasmids. This finding is counter-intuitive and in marked contrast with standard polymer systems (*37*) in which larger polymer sizes correlate with slower diffusion. This speed up is also observed in Differential Dynamic Microscopy experiments performed on entangled plasmids with different supercoiling degrees. Finally, we use sophisticated techniques involving minimal surface construction and primitive path analysis to quantify the abundance of threadings and entanglements between plasmids in solution and discover that larger supercoiling decreases both of these topological constraints, in turn explaining the enhanced mobility.

**Figure 1:**
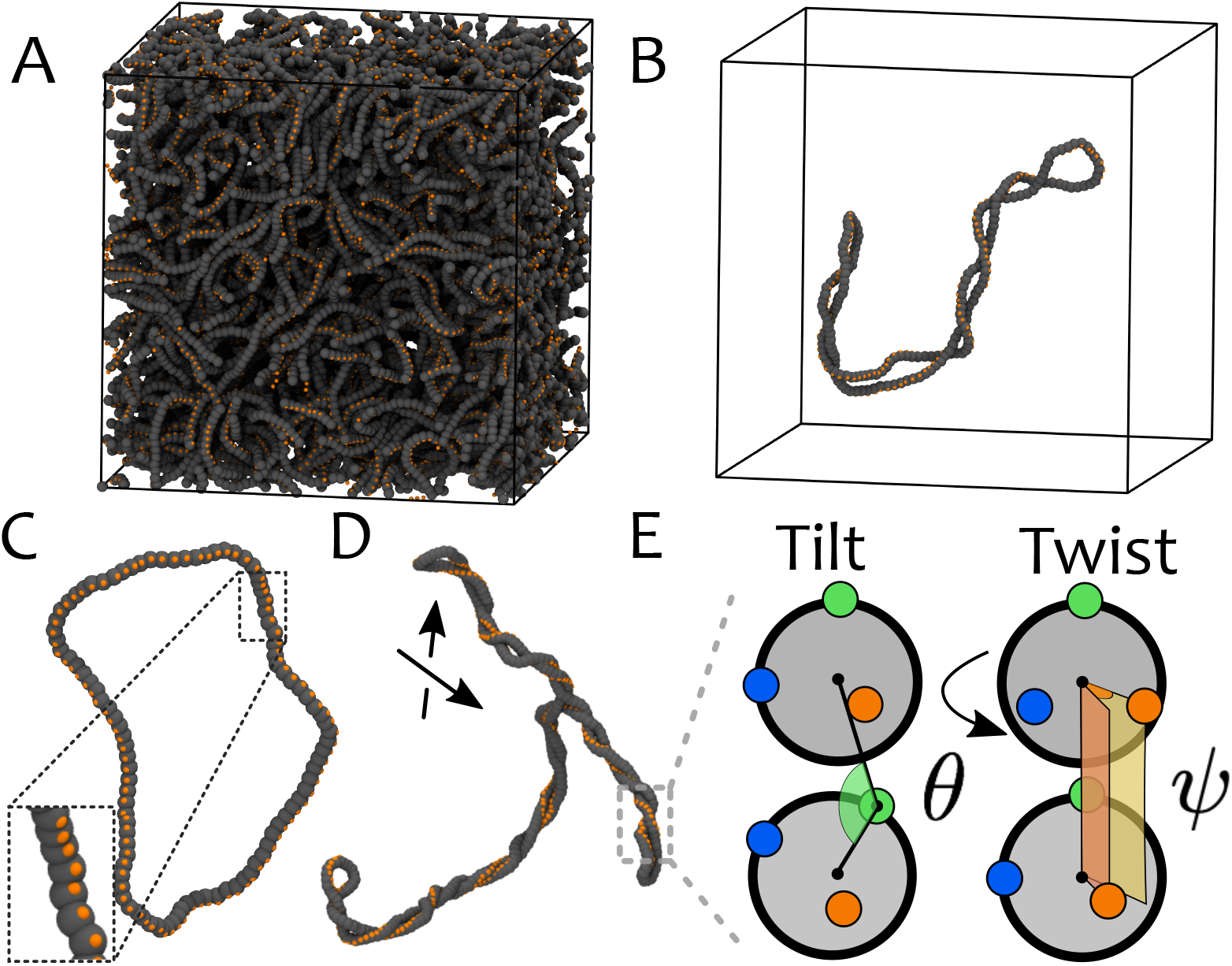
**A** Snapshot of simulation of entangled plasmids with length *L* = 200*σ_b_* ≃ 1.47 kbp and *σ* = 0.04. **B** A single plasmid taken from **A**. **C,D** Snapshots of plasmids with (**C**) *σ* = 0, *L* = 100*σ_b_* ≃ 750 bp and (**D**) *σ* = 0.06, *L* = 400*σ_b_* ≃ 3 kbp. Backbone beads are shown in grey, one set of patches are shown in orange. The other patches are not shown for clarity. **E** Sketch of tilt *θ* and twist *ψ* between consecutive beads (another angle *ψ* is set between blue patches, not shown). The tilt angle *θ* is subject to a stiff potential with equilibrium *θ*_0_ = *π* to maintain the frame co-planar and aligned with the backbone.

We argue that our results will be key to enabling the design of complex fluids with rheology that can be precisely tuned using a combination of DNA length, concentration, topology and supercoiling. Beyond providing blueprints for realising the next-generation of biomimetic DNA-based materials, our results can also shed light into the dynamics of DNA *in vivo*.

## Results and Discussion

### Computational Model for DNA plasmids

DNA is represented as a twistable elastic chain (*34, 38*) made of beads of size *σ_b_* = 2.5 nm = 7.35 bp connected by finitely-extensible springs and interacting via a purely repulsive Lennard-Jones potential to avoid spontaneous chain-crossing (*39*) (see Fig. 1). In addition to these potentials, a bending stiffness of *l_p_* = 50 nm (*7*) is set via a Kratky-Porod term and two torsional springs (dihedrals) constrain the relative rotation of consecutive beads, *ψ*, at a user-defined value *ψ*_0_. The torsional angle between consecutive beads *ψ* is determined by decorating each bead with three patches which provides a reference frame running along the DNA backbone. We finally impose a stiff harmonic spring to constrain the tilt angle *θ* = *π* so to align the frame with the backbone, i.e. along its local tangent (see Fig. 1D). The simulations are performed at fixed monomer density 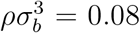 (corresponding to a volume fraction *ϕ* = 4% and *ϕ*/*ϕ*\* ≃ 16 with *ϕ*\* = 0.26%) and by evolving the equation of motion for the beads coupled to a heat bath in LAMMPS (*40*) (see Methods).

The user-defined angle *ψ*_0_ directly determines the thermodynamically preferred pitch of the twistable chains as *p* = 2*π*/*ψ*_0_ and, in turn, this fixes the preferred linking number to *Lk* = *M/p*, where *M* is the number of beads in the plasmid. The twist is enforced by an harmonic potential with stiffness *κ_t_* = 50*σ_b_* =125 nm comparable with the torsional persistence length of DNA. In this model, the relaxed linking number is *Lk*_0_ = 0 and so the degree of supercoiling *σ* ≡ Δ*Lk*/*M* = *Lk*/*M* = 1/*p*. The twist is set by initialising the patchy-polymer as a flat ribbon and by subsequently slowly increasing the stiffness of the potential associated with the twist degree of freedom. Ultimately, by imposing the angle *ψ*_0_ one can achieve the desired *σ* (which may be zero, if *ψ*_0_ = 0 or *p* = ∞). It should be noted that we will also consider non-torsionally constrained plasmids in which the torsional stiffness is set to *κ_t_* = 0 mimicking nicked, or enzymatically relaxed, circular plasmids. Finally, we recall that for supercoiled circular DNA, the exchange of local torsion (twist *Tw*) into bending (writhe *Wr*) must obey the White-Fuller-Calugareanu (WFC) (*41–43*) theorem, i.e. *Lk* = *Tw* + *Wr*, thus conserving the linking number *Lk* (and thus the supercoiling *σ* = Δ*Lk*/*M*) between the two DNA single strands (Fig. 1B-D). Notice that our polymer model is symmetric with respect to supercoiling and therefore we will refer to *σ* without specifying its sign.

### Supercoiling Increases the Average Size of DNA Plasmids in Entangled Conditions

The conformational properties of polymers in solution are typically studied in terms of the gyration tensor

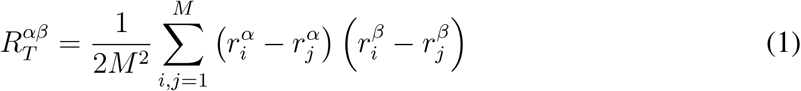

where 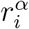 denotes the coordinate *α* of the position of bead *i*. The (square) radius of gyration is then defined as the trace, 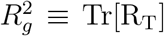. Interestingly, we find that the time and ensemble average of 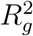 scales as 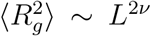, with metric exponents *ν* ≃ 1/2 and *ν* ≃ 3/5 for relaxed and highly supercoiled plasmids, respectively (for *M* ≥ 200, see Fig. 2A and Fig. S1 in SI). These two regimes are compatible with the ones for ideal (*ν* = 1/2) and self-avoiding (*ν* ≃ 0.588) random walks. This finding suggests that relaxed plasmids in entangled solutions (*ν* = 1/2) assume conformations similar to the ones of standard flexible, albeit short, ring polymers (*44*). At the same time, for larger values of *σ*, the self-interactions driven by writhing (see Fig. 1B,C) are no longer screened by the neighbours and we thus observe a larger metric exponent *ν* compatible with a self-avoiding walk.

**Figure 2:**
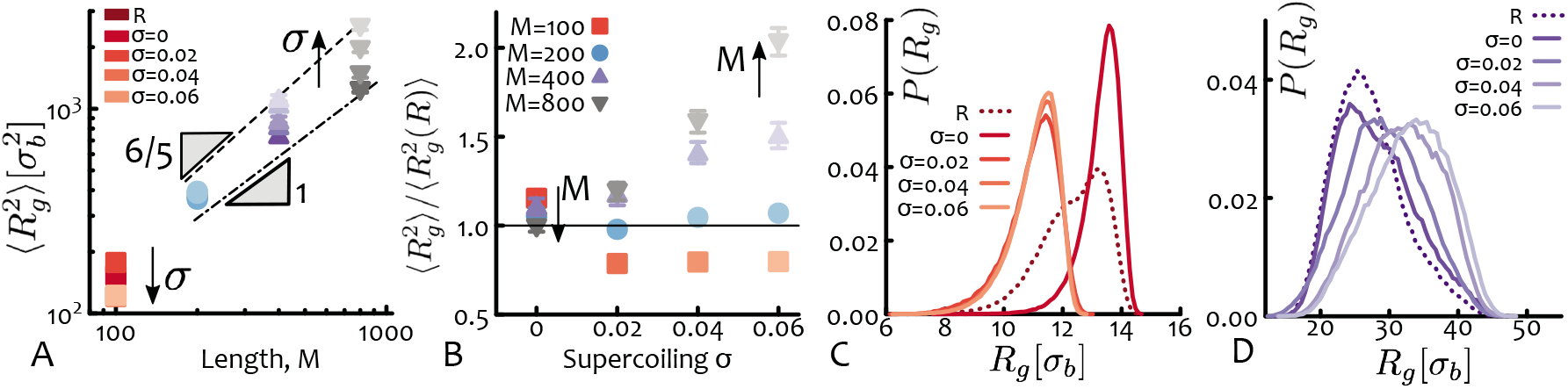
Supercoiling Increases Plasmids Size in Entangled Conditions. **A-B** Radius of gyrations *R_g_* plotted against (**A**) contour length *M* and (**B**) supercoiling *σ*. Notice that for short lengths *M* = 100, increasing *σ* induces a collapse of the plasmids whereas for longer lengths it drives swelling. The scaling of *R_g_* as a function of plasmid length is compatible with those of ideal and self-avoiding walks for relaxed and highly supercoiled plasmids, respectively. **C** The distribution of *R_g_* for *M* = 100 is weakly bimodal showing that plasmids can be in either an “open” or a “collapsed” state. Setting a supercoiling *σ* = 0 stabilises the open state whereas *σ* > 0 induces writhing and collapse. **D** For longer plasmids (*M* = 400) larger supercoiling *σ* broadens the distribution and drives enlarges the average size. The unit of length is *σ_b_* = 2.5 nm.

The effect of supercoiling on the average size of plasmids can be better appreciated in Fig. 2B where we show the (squared) radius of gyration rescaled by its value for relaxed plasmids and plotted against supercoiling. One can readily notice that, importantly, for long plasmids (e.g. *M* ≥ 400 ≃ 3 kb) the greater the supercoiling the *monotonically* larger their typical size. We highlight that this behaviour is highly counter-intuitive as one expects that supercoiling induces the compaction of a plasmid, as indeed is found computationally (*35*). At the same time, supercoiled plasmids travel faster than their relaxed counterparts in dilute conditions (*8*), e.g. in gel electrophoresis (*36*). Additionally, supercoiling is often associated with the packaging of the bacterial genome (*45, 46*) and with organisation into topological domains in eukaryotes (*14, 47, 48*). On the contrary, here we observe a monotonic increase of *R_g_* with supercoiling that is in marked contrast with the overall shrinking seen in dilute conditions (*35*) (this shrinking is also recapitulated by our model in dilute conditions, see Fig. S1 in SI).

We argue that this stark difference is due to inter-chain effects and the global topological invariance of the system. Indeed, while supercoiled plasmids may want to reduce their overall size, they also must remain topologically unlinked from the neighbours. In turn, the competition between this global topological constraint and the torsional and bending rigidities appears to favour swelling of long molecules (*L* > 200*σ* ≃ 1.5 kbp) but still drives the collapse of short ones (Fig. 2B).

For the shortest plasmids considered here (*M* = 100 ≃ 730bp), we observe an interesting exception to the behaviour described above whereby the typical size is non-monotonic for increasing supercoiling levels. More specifically, for *σ* = 0 we find that the conformations are typically larger than the relaxed ones, but they suddenly become more collapsed for *σ* > 0 (Fig. 2B). [Notice that with *σ* = 0 we mean plasmids that are intact and torsionally constrained to have linking number deficit equal to zero. These are different from relaxed (nicked) plasmids that are not torsionally constrained as the latter do not need to obey the WFC theorem; we denote them with ‘R’ throughout]. To investigate this behaviour further we examined the distributions of radius of gyration and noticed that relaxed short plasmids display a weakly bimodal distribution that is not found in larger plasmids (Fig. 2C,D). This bimodal distribution reflects the fact that short relaxed plasmids can be found in two typical conformational states: either open (large *R_g_*) or more collapsed (small *R_g_*) and imposing a certain supercoiling level appears to lock the molecules in one of the two states. In particular, since the conformational space of non-nicked plasmids must satisfy the WFC topological conservation law, zero supercoiling (*Lk* = *σ* = 0) hinders the writhing of the plasmid because it would be energetically too costly for them to writhe multiple times with opposite sign to achieve a null global writhe, given their short length (*L*/*l_p_* = 5). This entails that short plasmids with *σ* = 0 are locked into open, not self-entangled conformations. On the contrary, for *σ* > 0, the imposed writhing induces a conformational collapse, akin to a sharp buckling transition (*49*).

We note that the stable open state at *σ* = 0 for short plasmids is similar to the one computationally observed in dense solutions of semiflexible rings (*50*). These systems are expected to give rise to exotic columnar phases which would be thus intriguing to investigate in the context of dense solutions of short non-nicked plasmids with *σ* = 0.

We finally stress once more that the monotonic increase observed for long plasmids of their typical size with supercoiling is neither expected nor trivial and is in marked contrast with the overall shrinking behaviour found in the literature for long dilute supercoiled plasmids (*35*). Since the monomer concentration is constant for all the system studied, one would naïvely expect that these solutions are more entangled as *c*/*c*\* increases (the critical entanglement concentration *c*\* scales as 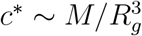). In turn, this implies more inter-chain entanglements which add up to the already more numerous intra-chain entanglements due to the increase of self-crossings (writhing). As a consequence, we would also expect highly supercoiled long plasmids to have reduced mobility with respect to their relaxed counterparts because of the more numerous entanglements with both themselves and also neighbouring chains.

### Supercoiling Enhances DNA Mobility

We study the dynamics of entangled plasmids at different levels of supercoiling by computing the time- and ensemble-averaged mean squared displacement (TAMSD) of the centre of mass (CM) of the plasmids as *g*_3_(*t*) = 〈***r***_*CM,i*_(*t*+*t*_0_) – ***r***_*CM,i*_(*t*_0_)〉_*i,t*_0__ (other *g_i_* quantities are reported in Fig. S4 in SI). Curves for *g*_3_ are shown in Fig. 3A,B for different values of plasmid supercoiling and length. Interestingly, and at odds with the findings of the previous section, we find that higher values of *σ* yield faster mobility especially for longer plasmids.

**Figure 3:**
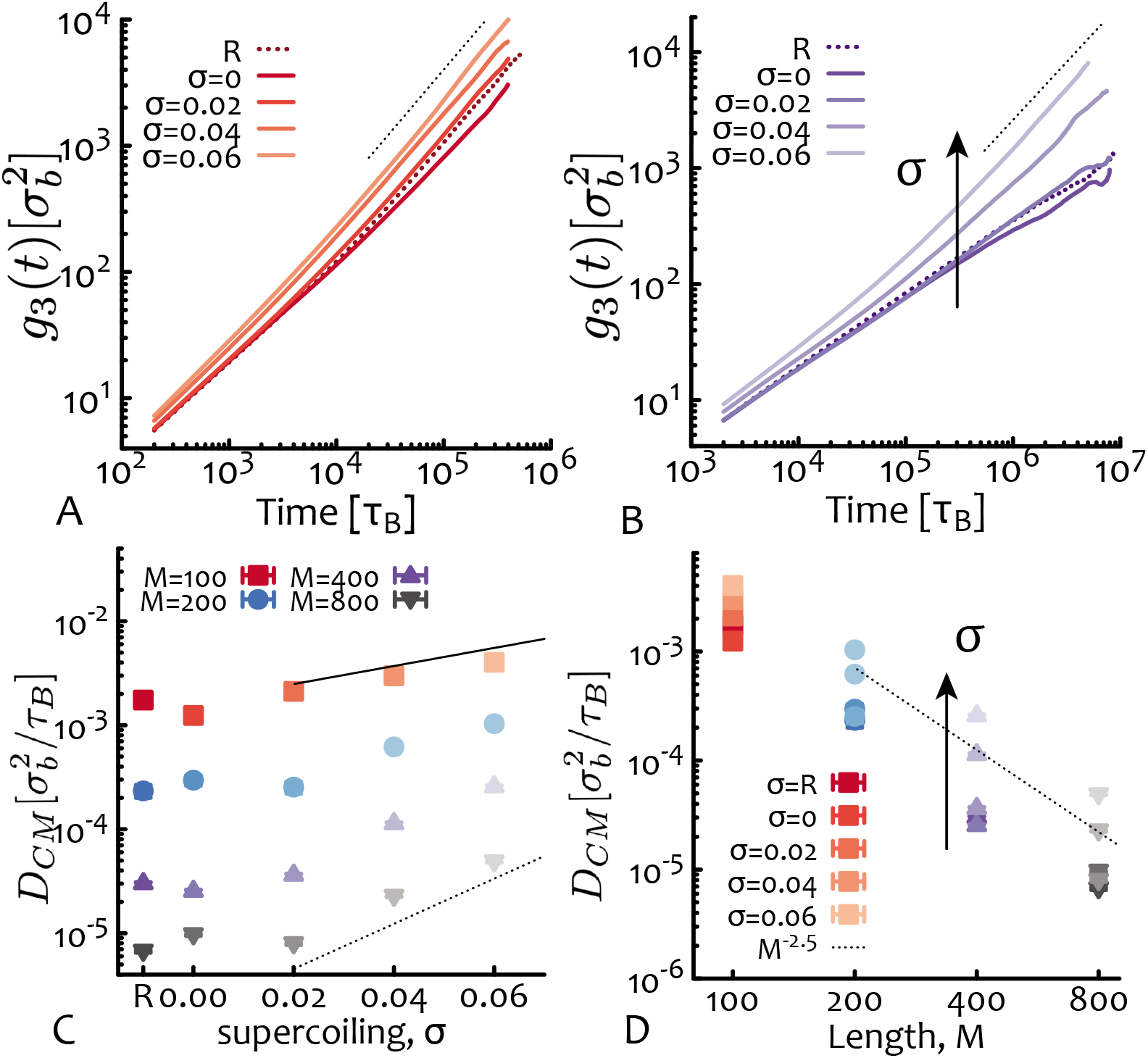
Supercoiling Enhances Plasmid Mobility. **A-B** Time-Averaged Mean Squared Displacement (TAMSD = *g*_3_) of the plasmids for (**A**) *M* = 100 ≃ 730 bp and (**B**) *M* = 400 ≃ 3 kbp. Dotted lines are linear functions of lagtime as a guide for the eye. **C-D** Diffusion coefficient of the centre of mass *D_CM_* = lim_*t*→∞_ *g*_3_(*t*)/6*t* against (**C**) supercoiling *σ* and (**D**) length *M*. In **C** exponentials ≃ exp (*σ*/0.05) (solid) and ≃ exp (*σ*/0.02) (dashed) are drawn as guide for the eye. In **D** a power law ≃ *M*^−2.5^ is fitted through the data for *σ* = 0.04. Error bars are comparable to symbol size. R = “relaxed”.

The diffusion coefficient of the centre of mass computed as *D_CM_* = lim_*t*→∞_ *g*_3_(*t*)/6*t* allows us to more precisely quantify how the mobility of the plasmids changes as a function of length and supercoiling. In particular, we find that while *D_CM_* attains a plateau at small *σ*, at larger supercoiling it increases exponentially (see Fig. 3C) albeit more simulations are needed to confirm this conjecture (see below for an argument supporting the exponentially faster mobility). Additionally, we find that the diffusion coefficient as a function of plasmid length scales as *D_CM_* ≃ *M*^−2.5^, compatible with the scaling of flexible and much longer ring polymers (*26*) (Fig. 3D). [Note that the solutions with *M* = 800 ≃ 6 kbp are not displaying a freely diffusive behaviour in spite of the fact that we ran them for more than 10^7^ Brownian times (see Tab. S1 in SI); in turn, *D_CM_* is overestimated as its calculation assumes free diffusion. In spite of this, values of *D_CM_* for *M* = 800 ≃ 6 kbp nicely follow the general trend of the other datasets (see Fig. 3C,D).]

It should be highlighted that the dynamical exponent found here is expected for longer and more flexible rings, and that for shorter flexible ring polymers (smaller *L*/*l_p_* and comparable with the rings studied here) the dynamical exponent is about 1.5 (*26*). We speculate that this more severe dependence on plasmid length may be due to the large persistence length of DNA (*l_p_* = 20*σ_b_* ≃ 50 nm) which can both, reduce the effective entanglement length (*27*) and also increase the number of threadings between plasmids (*51*). In agreement with this argument, entangled solutions of semi-flexible and stiff ring polymers vitrify in certain concentration regimes (*52*) implying that large stiffness and large concentrations may exacerbate threading constraints (*28, 53–55*).

### Differential Dynamic Microscopy of DNA plasmids confirm MD simulations

In order to experimentally validate the prediction that supercoiling enhances the mobility of plasmids in dense solutions we perform fluorescence microscopy experiments on 3 mg/ml solutions (corresponding to a volume fraction 0.4%) made of 6-kb plasmids. We label 0.001% of the molecules in solution and use Differential Dynamic Microscopy (DDM) to determine the diffusion coefficient from videos recorded on a custom fluorescence light-sheet microscope (*56*) (Fig. 4A). DDM, as compared to single-particle tracking, allows us the measure the dynamics of the diffusing molecules without having to resolve and track individual molecules over time – optimal for DNA of this size (*R_g_* < 100 nm). To pinpoint the role of supercoiling, we compare a solution of plasmids extracted from *E. Coli* in the stationary phase against the same solution pretreated with Topoisomerase I to relax the excess supercoiling (*57*) (see Methods).

**Figure 4:**
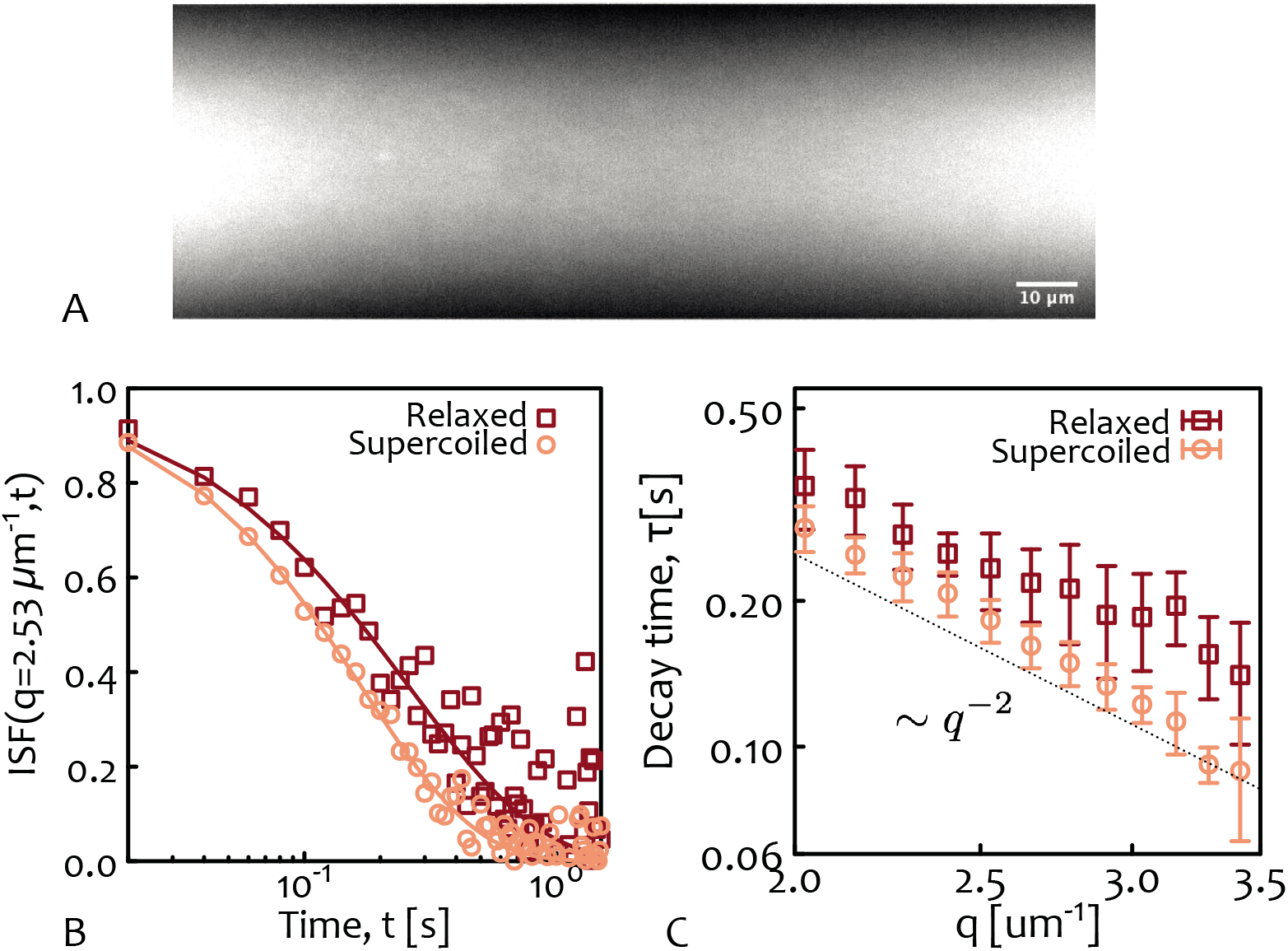
DDM of Entangled Plasmid DNA Confirms the Predictions from MD Simulations. **A** Snapshot from light-sheet microscopy showing fluorescent 5.9 kbp DNA plasmids (comparable with *M* = 800 is the MD simulations) at a concentration of 3 mg/ml concentration (*c*\* ≃ 0.6 mg/ml (*8*) and *c*/*c*\* ≃ 5). **B** Intermediate scattering function (ISF) obtained from DDM measurements. **C** Scaling of the ISF decay time with wave vector, showing that it scales as *q*^−2^. The fitted diffusion coefficients are *D* = 0.34(1) *μ*m^2^/*s* and *D* = 0.44(1) *μ*m^2^/*s* for relaxed and supercoiled plasmids respectively.

As one can notice (see Fig. 4B), the intermediate scattering function (ISF) shows a faster decay for supercoiled DNA compared to relaxed circular DNA, indicating faster dynamics. We fit each ISF with a stretched exponential *f*(*q, t*) = *exp*[−(*t*/*τ*)^*γ*^] using *γ* ≃ 0.9 −1 to determine the decay time *τ* as a function of *q* (Fig. 4C). As shown, the decay times are well fitted by a power law ≃ *q*^−2^ that we use to extract the diffusion coefficients via the relation *τ* = (2*Dq*^2^)^−1^. The resulting diffusion coefficients are *D* = 0.34(1) *μ*m^2^/*s* and *D* = 0.44(1) *μ*m^2^/*s* for relaxed and supercoiled solutions, respectively.

We should note that while our choice of plasmid length allows us to purify them without introducing substantial nicks (≃80% are without nicks and thus supercoiled), determining their precise supercoiling level is not straightforward. In vivo, supercoiling for plasmids in the stationary phase of cell growth (the phase at which we extract our plasmids) is ≃2% (*58, 59*). Thus, these results suggest that increasing supercoiling in solutions of entangled plasmids speeds them up and are thus in qualitative agreement with the simulations.

We should mention that while the experiments are at lower volume fraction with respect to simulations (when considering bare DNA), the buffering condition effectively thickens the diameter of DNA (*8*) thus rendering the precise comparison of experimental and simulated volume fractions difficult. We also note that due to the small size of the plasmids we are unable to accurately measure their size using single-molecule imaging. In turn, this renders the precise estimation of the overlap concentration also challenging (indirectly estimated to be about *c*\* ≃ 0.6 mg/ml (*8, 22*)). We are currently investigating alternative approaches, such as dynamic light scattering, so that in future work we can compare the intriguing predictions regarding the different sizes of supercoiled and relaxed circular DNA in dense solutions.

### Supercoiling Induces a Buckling Transition in Short Plasmids

The consequence of writhing on the plasmids conformations is not captured by *R_g_* alone (*60, 61*). Instead, it is informative to study shape descriptors which can be computed via the eigenvalues of the gyration tensor *R_T_* (which we denote as *a, b, c*, with *a* > *b* > *c* and 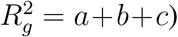). Typical shape descriptors are the asphericity (*60–62*) 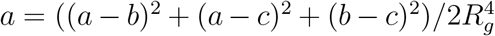 which quantifies the deviation from a perfectly spherical arrangement and the nature of asphericity quantified by either the prolateness (see Fig. S2 in SI) or the anisotropy 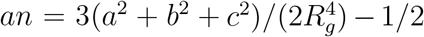 (shown in Fig. 5A,B). These shape descriptors reveal that for *M* = 100 ≃ 730 bp and *σ* = 0, plasmids are stabilised in an open, highly symmetric and oblate (M&M’s) state. Furthermore, they reveal that these short plasmids undergo a buckling transition to a closed, asymmetric and prolate (rugby ball) shape for *σ* > 0. The sharp first-order-like buckling transition (see Fig. 5A and SI) is weakened for larger contour lengths (see Fig. 5B), as self-writhing is energetically allowed even for *σ* = 0 (negative and positive self-crossings must cancel each other to satisfy the WFC conservation law). At the same time, both short and long plasmids display a general increase in asphericity, prolateness and anisotropy with increasing supercoiling, strongly suggesting that the plasmids assume elongated and double-folded conformations (see Fig. S2 in SI).

**Figure 5:**
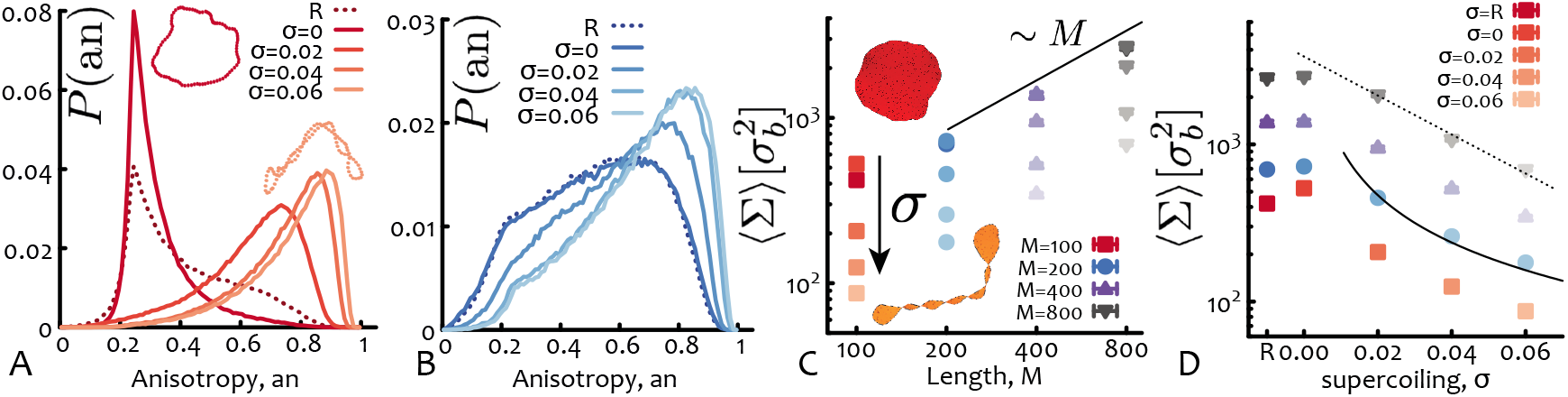
Supercoiling Induces Buckling in Short Plasmids and Reduces the Threadable Area. **A** The anisotropy shape descriptor (an, see text) for short plasmids *M* = 100 ≃ 730 bp displays a sharp buckling transition between an open and roughly symmetric state for *σ* = 0 and a collapsed and anisotropic one for *σ* > 0. In inset, two examples of conformations are shown. **B** For longer plasmids (*M* ≥ 200 ≃ 1.5 kbp) supercoiling shifts the anistropy to larger values indicating a smoother transition to more prolate conformations. **C** Scaling of the average minimal surface size 〈Σ〉 as a function of plasmids length (solid line shows the linear scaling). In inset, two examples of surfaces for *M* = 100 ≃ 730 bp are shown. **D** The size of the minimal surface area monotonically decreases with supercoiling (with the exception of short *M* ≤ 200 ≃ 1.5 kbp plasmids). The solid and dashed lines scale as 1/*σ* and *e*^−*σ*/0.035^, respectively, and are drawn as a guide for the eye. R = “relaxed”. The unit of length is *σ_b_* = 2.5 nm. The error bars, typically smaller than the symbol size represent the error of the mean area.

### Supercoiling Decreases the Spanning Minimal Surface

It is natural to associate the open-oblate/closed-prolate conformations assumed by DNA plasmids to a larger/smaller (minimal) spanning area, respectively (*63*). The size of this area may be relevant for the dynamics because it could be “threaded” by neighbouring plasmids hence hindering the dynamics (*28, 53, 54, 64*). To quantify this in more detail we calculated the minimal surface using the algorithm used in Ref. (*54, 63, 65*) for relaxed ring polymers. We found that the minimal area grows linearly with the plasmids’ contour, as expected (*54*) (Fig. 5C). Importantly, we also observed that it overall decreased with supercoiling with the notable exception of short *M* ≤ 200 ≃ 1.5 kbp plasmids, for which there is a small increase for *σ* = 0 with respect to the relaxed case, again confirming the topological locking in open conformations (Fig. 2A).

A crude way to estimate the decrease in “threadable” area of a plasmid is via recursive bisections of a perfect circle into several connected smaller circles joined at a vertex mimicking writhe-induced self-crossing. Each times a circle is split into two smaller ones the new radii are *R*′ ≃ *R*/2 and thus *n* circles (with *n* − 1 self-crossings) have radii *R*′ = *R*/*n* yielding an overall spanning surface ≃ *nπ*(*R/n*)^2^ ~ 1/*n* ~ 1/*σ*. This crude estimation is in good agreement with the behaviour of the minimal surface albeit we cannot rule out other functional forms (for instance exponential, see Fig. 5D). [Note that the so-called magnetic moment and radius (*66*) give similar results albeit different scaling (see Fig. S3 in SI)].

### Supercoiling Reduces Threadings

Motivated by the observation that the minimal surface – or “threadable area” – decreases with supercoiling, we decided to more precisely quantify the number of threadings per plasmid for different levels of supercoiling. To this end we identify a plasmid to be “passively threaded” by another when the minimal surface of the former is intersected by the contour of the latter (at least twice, as they are topologically unlinked) (*54*) (Fig. 6A). As shown in Fig. 6B, the average number of threadings per plasmid 〈*n_t_*〉 appears to decrease exponentially with supercoiling and to mirror the behaviour of 〈Σ〉. [As for the minimal surface, a notable exception to this general trend is the case of short plasmids (*M* = 100) for which we find that 〈*n_t_*〉 is statistically larger for *σ* = 0 than for relaxed plasmids, for the reason of “topological locking” that we explained above.]

**Figure 6:**
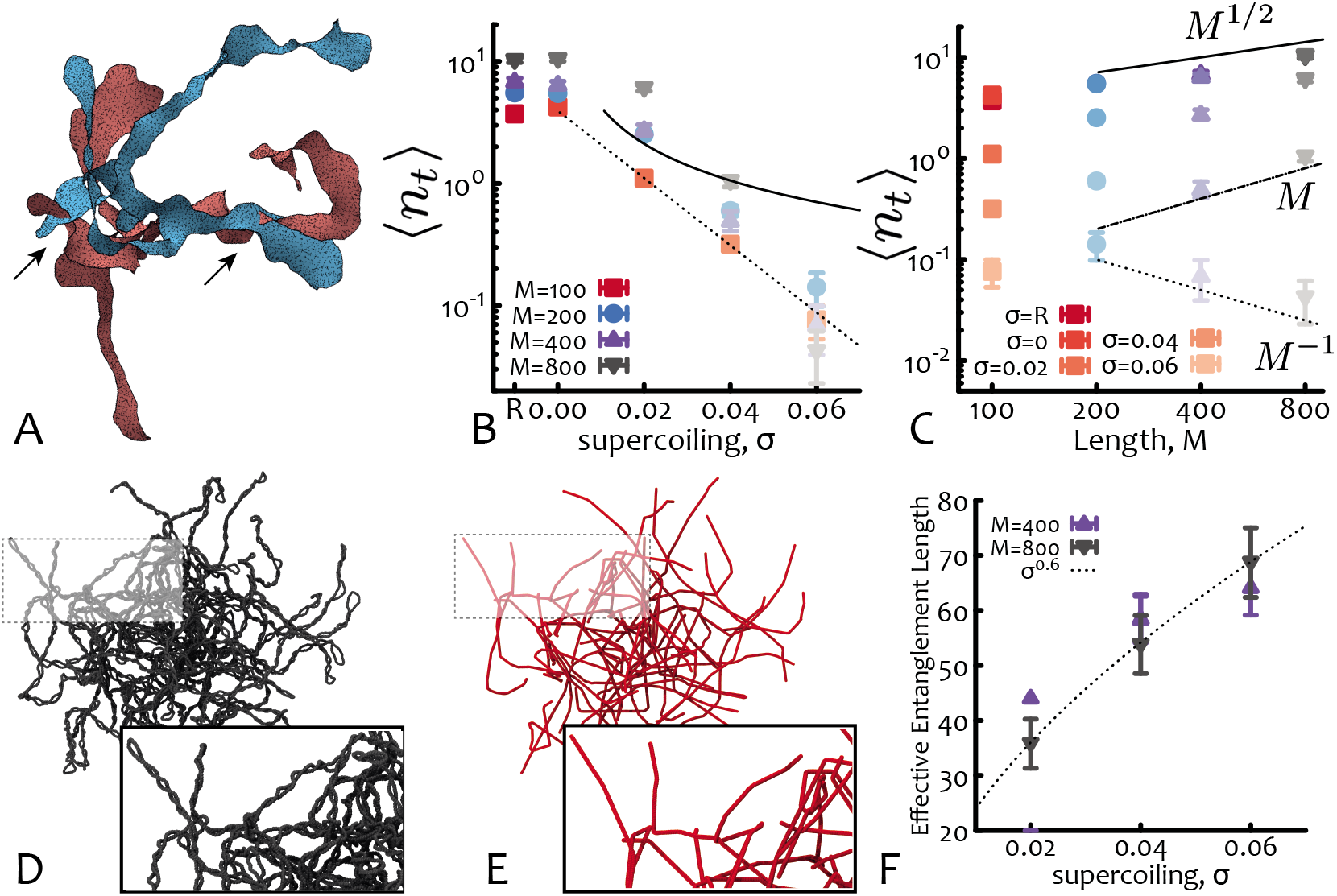
Supercoiling Reduces Threadings and Entanglements. **A** Snapshot of two threading plasmids (relaxed, *M* = 800 ≃ 6 kbp) with minimal surfaces drawn and intersections highlighted by arrows. **B** Number of threadings per plasmid as a function of supercoiling (dashed=exponential, solid=1/*σ*). **C** Number of threadings per plasmid as a function of DNA length (dashed=1/*M*, solid=*M*^1/2^, dot-dashed=*M*). **D,E** Snapshots of the PPA analysis run on a system with plasmids *M* = 800 ≃ 6 kbp and *σ* = 0.06. **F** The effective entanglement length increases with supercoiling as *N_e_* ≃ *s*^0.6^ for *M* = 800 ≃ 6 kbp.

Based on these findings, we can also advance an argument as for why the diffusion co-efficient of plasmids should increase exponentially with supercoiling: since the dynamics of polymers with threadings – for instance entangled rings (*64, 67*), melts of tadpole-shaped polymers (*24, 68*) or compressed long plasmids (*69*) – slows down exponentially with the number of threadings (*24*), the fact that increasing supercoiling reduces the threadable area and the number of threadings entails an exponential increase in diffusion. We thus expect the dynamics of highly supercoiled (threading-poor) plasmids to be exponentially faster than their relaxed (threading-rich) counterparts as seen in Fig. 3C.

Curiously, in the case of short plasmids in which setting *σ* = 0 increases the threadable area and also the number of threadings we also find a slower dynamics, in full agreement with our argument.

### Supercoiling Reduces Entanglements

The shape descriptors studied above suggest that long plasmids assume prolate double-folded conformations, but it remains unclear whether the conformations are simply plectonemic (linear-like) or more branched into comb, star or tree-like structures (*70*). We thus computed the local absolute writhe along the contour length, *W*(*s*), from which the number and location of plectonemic tips can be extracted as arg max *W*(*s*) (*71, 72*) (see Methods). This calculation reveals that most of the conformations with *σ* ≥ 0.04 have 2 tips and so are mainly linear-like plectonemic conformations (see Fig. S5 in SI). [For smaller supercoiling it is difficult to unambiguously distinguish tips from other regions of large curvature.]

In light of this finding another apparent controversy arises. Indeed, arguably, linear chains half the length as their ring counterparts are expected to diffuse slower than the rings due to reptation relaxation induced by ordinary entanglements (assuming that the entanglement length is the same for the two systems) (*26*); instead, we observe the opposite trend. To explain this result we adapted the primitive path analysis (PPA) method (*73*) (see Fig. 6C,D and Methods) and determined an effective entanglement length *N_e_* for highly supercoiled plasmids leveraging the fact that the tips of linear-like or branched conformations represent effective termini that can be pinned down. [Note that the PPA method typically fails for standard relaxed ring polymers because there are no ends to pin]. We find that *N_e_* grows with *σ* (Fig. 6F), suggesting that the larger the supercoiling the less entangled the plasmids. We argue that this effective reduction in entanglements (opposite to what one would naively expect considering 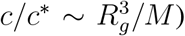) is due to the fact that supercoiling increases the local concentration of intra-chain beads (*74*) in turn inflating the effective tube surrounding any one plasmid.

Notice that for short plasmids the PPA method cannot identify an entanglement length, confirming that these are very poorly entangled in the standard sense. As such, their dynamics is mostly determined by threadings, which are abundant also in short plasmids (see Fig. 6B,C).

#### Conclusions

In this work we have studied the dynamics of entangled solutions of DNA plasmids to understand how supercoiling can be leveraged to tune the rheology of dense DNA solutions orthogonally to other traditional methods, such as varying length or concentration. We have discovered that, contrary to what is typically assumed, the size of long plasmids increases with supercoiling when in entangled solutions. Surprisingly, we find this swelling is mirrored by an enhanced mobility. Our predictions are supported by experiments which show that the diffusion coefficient of entangled intact and supercoiled (*σ* ≃ 0.02) plasmids is larger than that of relaxed ones, i.e. with *σ* ≃ 0. We discovered that this enhanced mobility is due to severely asymmetric conformations which greatly reduce the number of threadings. In parallel, entanglements are also reduced as supercoiling increases the local concentration of intra-chain contacts thus effectively inflating the tube formed by neighbouring chains. We thus discovered that the unexpected enhanced diffusivity of entangled supercoiled DNA is due to a combination of reduced entanglements and, in particular, threadings. We note that threadings are abundant also in short plasmids that are poorly entangled in the standard sense and thus play a major role in determining their dynamics.

We conjecture that beyond the range of lengths studied in this work (0.7 – 6 kbp), supercoiled plasmids in entangled solutions may display branched architectures, triggering the need of arm retraction or plectoneme diffusion/hopping relaxation mechanisms; as these are notoriously slow processes, we predict a re-entrant slowing down of the plasmid diffusion. This non-monotonic dependence of DNA mobility on supercoiling would allow even richer control of the rheological properties.

In summary, our results define a clear route for the topological tuning of the rheology of DNA-based complex fluids that employs supercoiling as a mean to control DNA mobility. We note that the fact that supercoiling regulates the number of threadings per plasmids can also be leveraged in polydisperse systems or in blends of linear and supercoiled DNA or other biopolymer composites, where threading of rings by the linear contaminants is key to determine the stress relaxation of materials (*25*).

In the future, it would be interesting to further investigate either longer plasmids computationally or plasmids with selected level of supercoiling in experiments. Albeit difficult, this may be feasible using caesium chloride gradient separation techniques (*75*). Ultimately, understanding how DNA topology and supercoiling affect the dynamics and conformational properties of plasmids in *entangled* or *crowded* conditions may not only reveal novel pathways to achieve fine tuning of the rheology of complex biopolymer fluids but also shed light on the role of supercoiling in vivo.

## Material and Methods

### Molecular Dynamics

Each bead in our simulation is evolved through the Langevin equation 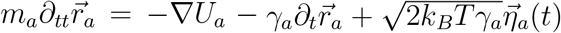, where *m_a_* and *γ_a_* are the mass and the friction coefficient of bead *a*, and 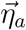 is its stochastic noise vector satisfying the fluctuation-dissipation theorem. *U* is the sum of the energy fields (see SI). The simulations are performed in LAMMPS (*40*) with *m* = *γ* = *k_B_* = *T* = 1 and using a velocity-Verlet algorithm with integration time step Δ*t* = 0.002 *τ_B_*, where *τ_B_* = *γσ*^2^/*k_B_T* is the Brownian time.

### Branching Analysis

Following Refs. (*71, 76*), we compute the absolute writhe of a segment of a plasmid as 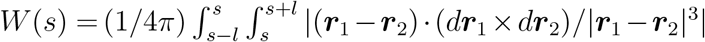 with window *l* = 50 beads. This calculation yields a function *W*(*s*) whose maxima represent regions of high local writhe and can identify tips of plectonemes. In addition to being a local maximum, we require that *W*(*s*) > 0.35 to avoid false positives. See SI for more details.

### Primitive Path Analysis

Following Ref. (*73*), we fix certain polymer segments in space, turn intra-chain repulsive interactions off and keep inter-chain interactions on. We then run simulations at low temperature 0.01 to find a ground state. The resulting chain conformations (primitive paths) are made of straight segments connected by sharp kinks due to entanglements. The entanglement length is then given by 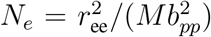, where *r*_ee_ is the mean endpoint distance, *M* is the number of monomers between the fixed points and *b_pp_* is the mean bond-length of the primitive path. We adapt the classical PPA for plasmids by fixing the tips of all detected plectonemes instead of the end points of linear chains (see SI).

### DNA preparation

Double-stranded 5.9 kbp DNA plasmids are replicated in E. coli, collected at the onset of stationary phase, before being extracted and purified using our previously described protocols (*8*). Following purification the DNA solution is ~80% supercoiled and ~20% relaxed circular, as determined from gel electrophoresis (see SI, Fig. S6). To produce concentrated solutions of relaxed circular DNA, Topoisomerase I (New England Biolabs) is used to convert the DNA topology from supercoiled to relaxed circular (*77*) (see SI, Fig. S6). Both supercoiled and relaxed circular DNA solutions are concentrated to 3 mg/ml using an Eppendorf Vacufuge 5301.

### Fluorescence Imaging

To visualize DNA diffusion in concentrated solutions, supercoiled or relaxed circular DNA is labeled with YOYO-1 dye (Thermo Fisher Scientific) at a 4:1 base pair:dye ratio (*78*), and added at a concentration of 0.045 *μ*g/ml to 3 mg/ml solutions of supercoiled or relaxed circular DNA described above. Glucose (0.9 mg/ml), glucose oxidase (0.86 mg/ml), and catalase (0.14 mg/ml) are added to inhibit photobleaching (*57, 79*). The DNA solutions are pipetted into capillary tubing that is index-matched to water and imaged using a custom-built light-sheet microscope with a 488 nm excitation laser, an excitation objective of 10x 0.25 numerical aperture (NA), an imaging objective of 20x 1.0 NA, and an Andor Zyla 4.2 CMOS camera. At least 4 sample videos are recorded at 50 frames per second for 2000 frames. The video dimensions are 256 x 768 pixels, which are then analyzed by examining regions of interest (ROI) of 256 x 256 pixels (50 x 50 *μ*m).

### DDM analysis

We follow methods previously described to investigate DNA diffusion using Differential Dynamic Microscopy (DDM) (*57, 80, 81*). Briefly, from each ROI we obtain the image structure function or DDM matrix *D*(*q*, Δ*t*), where *q* is the magnitude of the wave vector and Δ*t* is the lag time. To extract the transport dynamics of the diffusing DNA molecules, we fit the structure functions to *D*(*q*, Δ*t*) = *A*(*q*)[1 – *f*(*q*, Δ*t*)] + *B*(*q*), where *B* is a measure of the camera noise, *A* depends on the optical properties of both the sample and microscope, and *f*(*q*, Δ*t*) is the intermediate scattering function (ISF). Based on our previous studies of microspheres and DNA diffusing in crowded environments, we fit the ISFs to stretched exponentials of the form *f*(*q*, Δ*t*) = exp – (Δ*t*/*τ*(*q*))^*γ*(*q*)^, where *τ* is the characteristic decay time and *γ* is the stretching exponent, both of which depend on *q* (*57, 81–83*).

For normal free diffusion, one expects ISFs described by a simple exponential, i.e., *γ* =1, while our scattering functions are better fitted with stretching exponents between 0.9–1. Having extracted the decay times of density fluctuations *τ* over a range of spatial frequencies *q*, we fit the results to *τ* = (2*Dq*^2^)^−1^ to determine the diffusion coefficient, *D*, for the DNA plasmids.

## Acknowledgments

The authors would like to acknowledge the networking support by the “European Topology Interdisciplinary Action” (EUTOPIA) CA17139. This project has received funding from the European Union’s Horizon 2020 programme under grant agreement No. 731019 (EUSMI). DM acknowledges the computing time provided on the supercomputer JURECA at Jülich Supercomputing Centre and support by the Leverhulme Trust through an early career fellowship (ECF-2019-088). JS acknowledges the support from the Austrian Science Fund (FWF) through the Lise-Meitner Fellowship No. M 2470-N28. JS is grateful for the computational time at Vienna Scientific Cluster. Sample codes can be found at git.ecdf.ed.ac.uk/dmichiel/supercoiledplasmids.

## Supplementary materials

Materials and Methods

Supplementary Text

Figs. S1 to S5

Tables S1

